## Supplementary figures and images for "Enhancing Bone Regeneration and Osseointegration using rhPTH(1-34) and Dimeric ^R25C^PTH(1-34) in an Osteoporotic Beagle Model"

### suppl

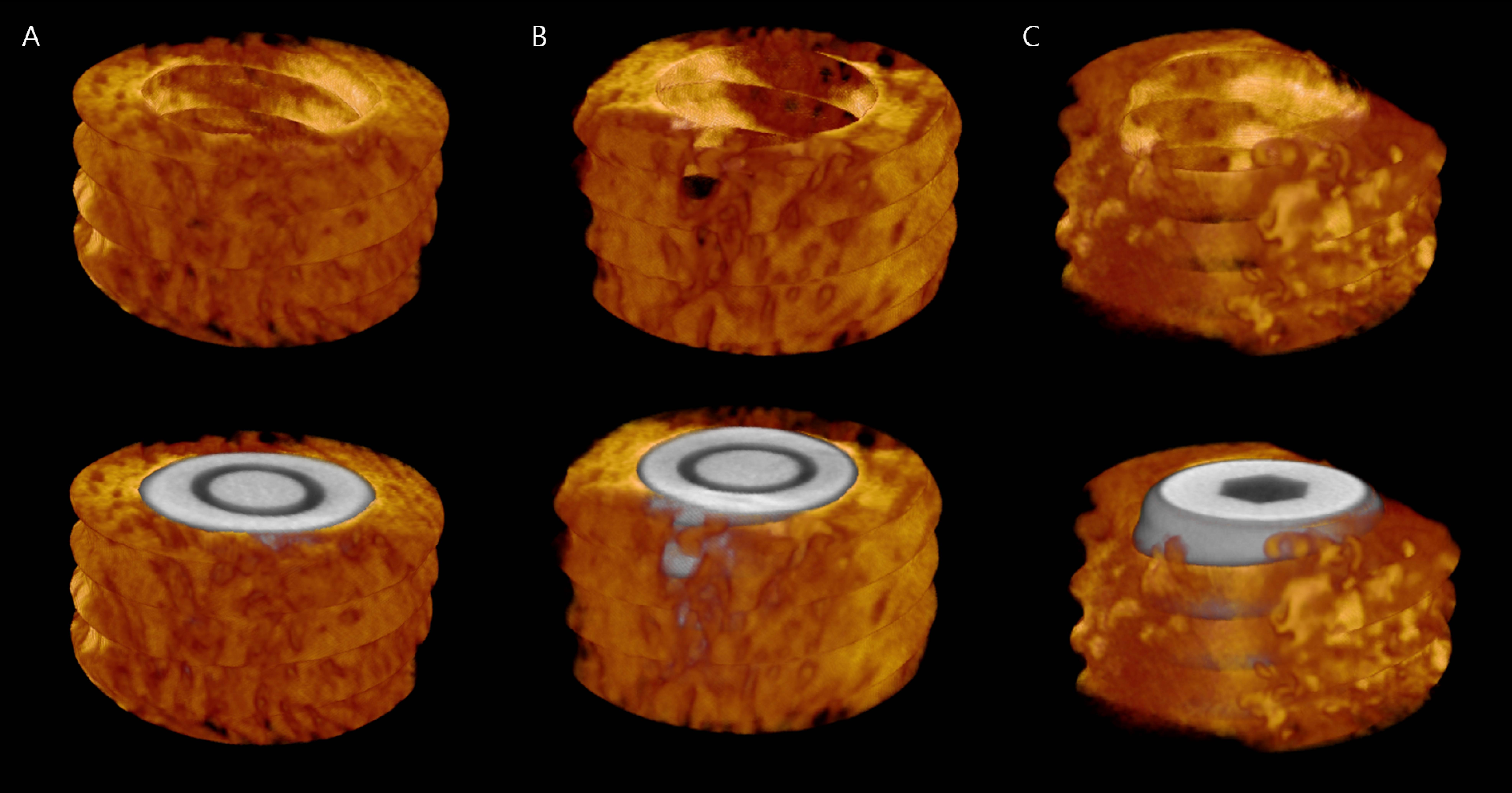
